# One read per cell per gene is optimal for single-cell RNA-Seq

**DOI:** 10.1101/389296

**Authors:** M. J. Zhang, V. Ntranos, D. Tse

## Abstract

An underlying question for virtually all single-cell RNA sequencing experiments is how to allocate the limited sequencing budget: deep sequencing of a few cells or shallow sequencing of many cells? A mathematical framework reveals that, for estimating many important gene properties, the optimal allocation is to sequence at the depth of one read per cell per gene. Interestingly, the corresponding optimal estimator is not the widely-used plugin estimator but one developed via empirical Bayes.

---

Single-cell RNA sequencing (scRNA-Seq) technologies have revolutionized biological research over the past few years by providing the tools to simultaneously interrogate the transcriptional states of thousands of cells in a single experiment. In contrast to bulk RNA-Seq which probes the *average* gene expression in a cell population, single-cell RNA-Seq has unlocked the potential of extracting higher-order information, granting us access to the underlying gene expression distribution. Indeed, this unprecedented look into population-level heterogeneity has been key in the success of scRNA-Seq leading up to new biological discoveries [1–2].

Although early single-cell RNA-Seq assays were labour intensive and initially constrained by the small number of cells that could be processed in a single experiment, recent technological advances have allowed hundreds of thousands of cells to be assayed in parallel [3], eliminating the otherwise prohibitive per cell cost overhead. From a sequencing budget perspective, however, this seemingly unconstrained increase in the number of cells available for scRNA-Seq introduces a practical limitation in the total number of reads that can be sequenced per cell. More reads can significantly reduce the effect of the technical noise in estimating the true transcriptional state of a given cell, while more cells can provide us with a broader view of the biological variability in the population. A natural experimental design question arises (Fig. 1a): how many cells should we choose to profile for a given study and at what sequencing depth?

**Figure 1.**
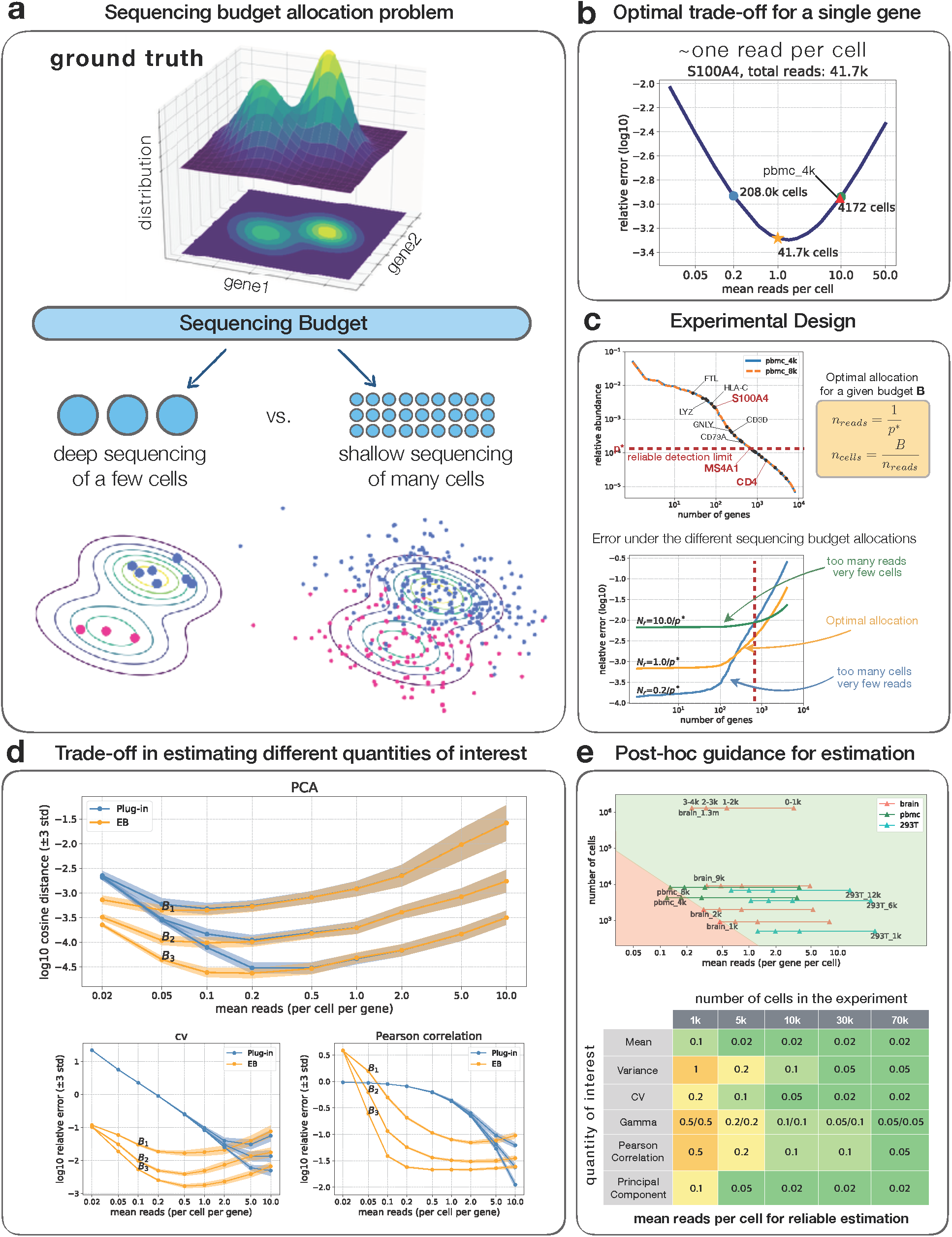
**a) Sequencing budget allocation problem.** Consider estimating the underlying gene distribution (top) from the noisy read counts obtained via sequencing (bottom). With a fixed number of reads to be sequenced, deep sequencing of a few cells accurately estimates each individual cell but lacks a coverage of the entire distribution (left), while a shallow sequencing of many cells covers the entire population but introduces a lot of noise (right). **b) Optimal trade-off**. The memory T-cell marker gene S100A4 has 41.7k reads in the pbmc_4k dataset. For estimating the underlying gamma distribution 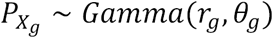, the relative error is plotted as a function of the sequencing depth, where the optimal error is obtained at the depth of one read per cell (orange star) and is 2 times smaller than that at the current depth of pbmc_4k (red triangle). **c) Experimental design**. To determine the sequencing depth for an experiment, first the relative gene expression level can be obtained via bulk RNA-Seq (top left). Second the researcher can select a set of genes of interest (i.e. some marker genes highlighted as black dots), of which the smallest relative expression level < (MS4A1) defines the reliable detection limit. Finally, the optimal sequencing depth is determined as *n*_*reads \**_ *\** = 1/*P*\* (top right). The errors under different trade-offs are visualized as a function of the genes ordered from the most expressed to the least (bottom). The optimal sequencing (orange) minimizes the worst-case error over all the genes of interest (left of the red dashed line), while both the deeper sequencing (green) and the shallower sequencing (blue) yield worse results. **d) Simulation of error under different budget allocation**. The top panel simulates the error for estimating the first principal direction using plug-in (blue) and EB (orange) respectively. Three budgets are considered *B*_1_ (0.6k/gene) < *B*_2_(3k) < *B*_3_(15k) and the depth (mean reads per cell per gene) ranges from 0.02 to 10. The optimal depth for EB is the same (∼0.1) for all three budgets, validating the theory that the optimal depth is independent of the budget. The cases for the coefficient of variation and the Pearson correlation (bottom) also show similar qualitative behaviours. **e) Post-hoc guidance for reliable estimation**. We visualized the top 4k genes of some popular datasets (top), where a triangle residing in the green region means the Pearson correlation of corresponding genes can be reliably estimated (relative error<10%). For example, we can reliably estimate the first 2k genes for the brain_1k dataset and all 4k genes for the brain_9k dataset. A more comprehensive result is summarized in the bottom table. For example, the first element (mean, 1k) shows that with 1k cells, a gene need to have at least 0.1 reads per cell for reliably estimating the mean.

The experimental design question has attracted a lot of attention in the literature [4–8], but as of now there has not been a clear answer. Several studies provide evidence that a relatively shallow sequencing depth is sufficient for common tasks such as cell type identification and principal component analysis (PCA) [9–11], while others recommend deeper sequencing for accurate gene expression estimation [12–15]. Despite the different recommendations, the approach to providing experimental design guidelines is shared among all: given a deeply sequenced dataset with a predefined number of cells, how much subsampling can a given method tolerate? An example of this conventional approach is also evident in the mathematical model used in [11] to study the effect of sequencing depth on PCA. Although practically relevant, this line of work does not provide a comprehensive solution to the underlying experimental design question because of three reasons: 1) the number of cells is *fixed* and implicitly assumed to be enough for the biological question at hand; 2) the deeply sequenced dataset is considered to be the ground truth; 3) the corresponding estimation method is chosen a *priori* and is tied to the experiment.

In this work, we propose a mathematical framework for single-cell RNA-Seq that fixes not the number of cells but the total sequencing budget, and disentangles the biological ground truth from both the sequencing experiment as well as the method used to estimate it. In particular, we consider the output of the sequencing experiment as a noisy measurement of the true underlying gene expression and evaluate our fundamental ability to recover the gene expression distribution using the *optimal* estimator. The two design parameters in our proposed framework are the total number of cells to be sequenced *n*_*cells*_ and the sequencing depth in terms of the total number of reads per cell *n*_*reads*_ both of which affect the optimal estimation error. Now, the experimental design trade-off becomes apparent when these two quantities are tied together under a total sequencing budget constraint *B* = *n*_*cells*_ × *n*_*reads*_ (Fig. 1a, sequencing budget allocation problem). More specifically, we consider a hierarchical model [16] to analyze the trade-off in the sequencing budget allocation problem (see Methods). At a high level, we assume an underlying gene expression distribution *P*_**X**_ that carries the biological information we are interested in and is independent of the sequencing process (Fig. 1a top). The cells in the experiment are therefore associated with gene expression levels **X**_1_,**X**_2_,… sampled from *P*_**X**_, while we can only observe the read counts **Y**_1_,**Y**_2_,… that are generated from the corresponding gene expression levels via sequencing (Fig. 1a bottom). In this context, it is clear that with many cells *n*_*cells*_ we can estimate the read count distribution *P*_**Y**_ accurately while with more reads per cell *n*_*reads*_ we can make sure that individual observations **Y**_1_,**Y**_2_,… are much closer to the ground truth expression levels **X**_1_,**X**_2_… of the cells. The optimal trade-off is then derived to reconcile the two.

For our main results, we focused on 3′-end sequencing technologies [17–19] and used the above framework to study the experimental design trade-off for estimating several important quantities of the underlying gene distribution, such as the coefficient of variation (cv) and the Pearson correlation (Methods). In the context of 3′-end sequencing, *P*_**X**_ naturally models the unknown high-dimensional distribution of mRNA abundances across cells while the read counts for the cells, **Y**_1_,**Y**_2_,… correspond to the number of unique molecular identifiers (UMIs [20]) observed via sequencing. Our main result states that the optimal budget allocation (i.e., the one that minimizes the estimation error) is achieved by maximizing the number of cells while making sure that at least ∼1 UMI per cell will be observed for the genes of biological interest to the experiment. Although our framework is non-parametric — in the sense that no particular prior is assumed for the underlying gene distribution *P*_**X**_, it is instructive to illustrate the result in the context of the widely-used negative binomial (NB) overdispersion model (Methods, Supp. Note 5.1). In this case, one would be interested in estimating the underlying gamma distribution *P*_**X**_ ∼ *Gamma*(**r**,**θ**) that gives rise to the NB-distributed counts after sequencing via Poisson sampling, effectively distinguishing the biological variability from the technical variability. As a demonstrating example, in Fig. 1b we consider the memory T-cell marker gene S100A4 to be of biological interest and evaluate the optimal trade-off for the total sequencing budget used to generate the 10x Genomics’ pbmc_4k dataset (4340 cells, Supp. Note 6.2); our analysis suggests that the optimal trade-off would have been attained by sequencing 10 times shallower using 10 times as many cells, reducing the error by two folds. The recommended sequencing depth depends on the genes under consideration. For example, the sequencing depth of pbmc_4k dataset is optimal when the B-cell marker gene MS4A1 is considered and it should be sequenced 4 times deeper with ¼ cells when the T-helper marker gene CD4 is considered (Fig. 1c top, Supp. Fig. 2a-b). The latter is arguably to have reached saturation for the 10x Genomics technology. Hence, the guidance there is to sequence until saturation (Methods).

**Figure 2.**
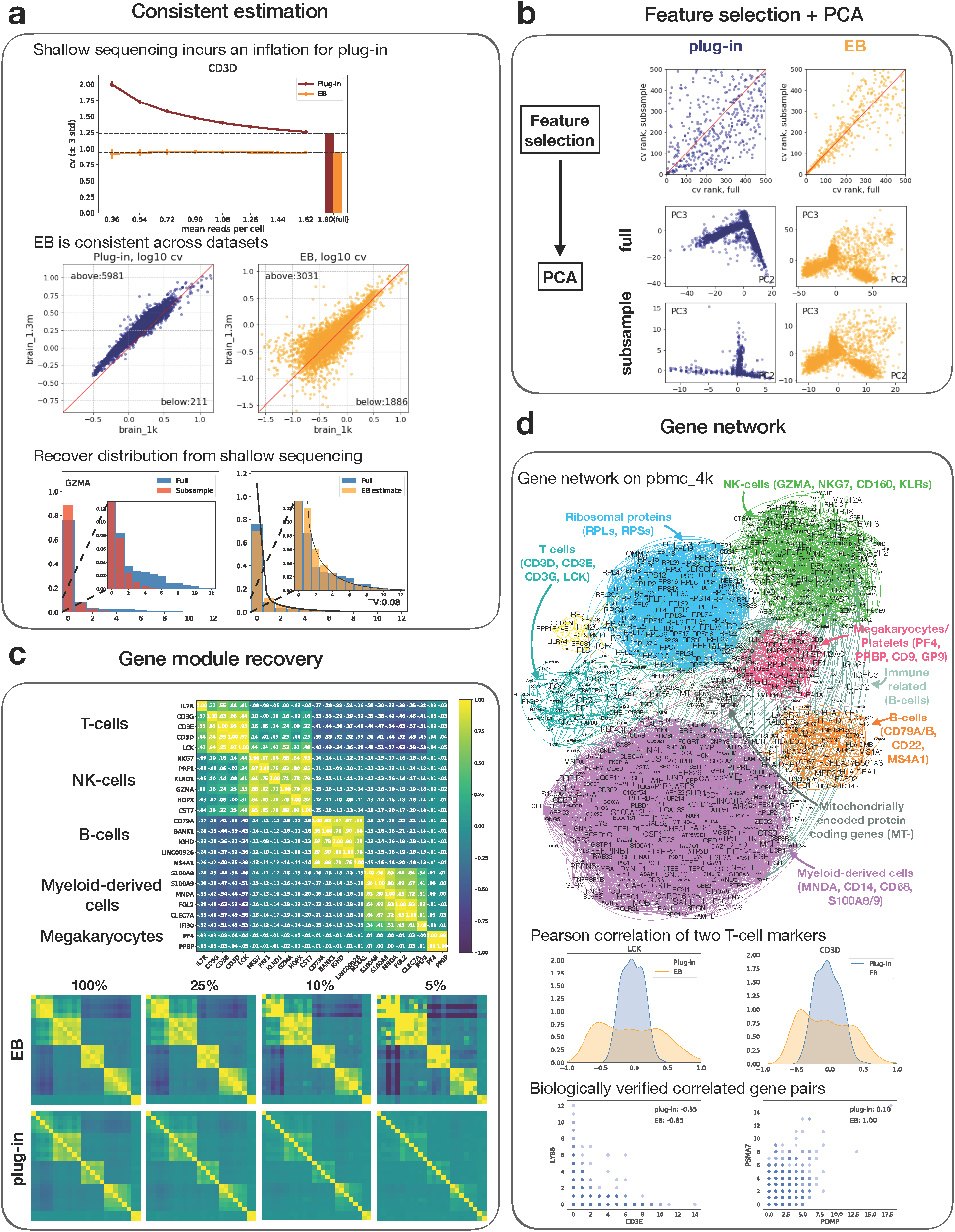
**a) Consistent estimation. Top:** For estimating the cv, the plug-in estimates get more inflated as the sequencing depth become shallower (from right to left along the x-axis) while the EB estimates are nonetheless consistent. **Middle**: brain_1k and brain_1.3m are both from E18 mice brain. Hence a similar estimated cv is expected for each gene. This is indeed the case for EB (right), which is adaptive to different sequencing depths. However, since brain_1k is twice deeper than brain_1.3m, a bias is introduced for plug-in that most points are above the 45-degree line (red). **Bottom**: EB recovers the distribution for the gene GZMA from a data that is subsampled to be 5times shallower (left). The result shows a good estimation of the zero proportion as well as the tail shape, thus incurring a small total variation error (TV) (right). **b) Feature selection and PCA**. The task is to first select features (genes) based on cv and then perform PCA on the selected features. The results on the full data (pbmc_4k) and a subsampled are compared, where the subsampled data is 3 times shallower. Both the cv ranks (top) and the PCA plots (bottom) are more consistent for EB between the full data and the subsampled data. **c) Gene module recovery**. The EB estimated Pearson correlation for the marker genes in pbmc_4k are visualized and are sorted according to different cell populations (top). The clear block-diagonal structure implies that EB can well capture the gene functional groups. As a comparison, the plug-in estimator also recovers those modules but with a weaker contrast (bottom left most). A subsample experiment is further conducted to investigate how well the estimators can recover the modules from the shallow sequencing data. EB can recover the module at the depth of 10% while for plug-in, the first block (T-cells) is blurred at the depth of 25% and the entire structure vanishes at the depth of 10%. **d) Gene network**. **Top**: a gene network based on the EB estimated Pearson correlation using the pbmc_4k dataset. Most gene modules are found to correspond to either important cell populations or cell functions, including T-cells, B-cells, NK-cells, myeloid-derived cells, megakaryocytes/platelets, ribosomal protein genes, and mitochondrially encoded protein coding genes. **Middle**: the estimated Pearson correlations of all genes with LCK and CD3D respectively, where the two genes are known to be enriched in T-cells. There are three modes for the EB estimated values, where the positive mode, the zero mode, and the negative mode correspond to genes in the same module, different modules, and irrelevant genes respectively. The plug-in estimated values are nonetheless much closer to zero even for the truly correlated ones, indicating an artificial shrinkage of the estimated values. **Bottom**: two instances where EB significantly differs from plug-in. The axes represent read counts and the dots show color-coded histogram over the cells. Both pairs of genes are biologically validated (Methods).

As the example indicates, an important aspect of our framework is to allow flexible experimental design at a single-gene resolution. The researcher can thus design the experiment based on the mean expression level of a set of important genes related to the biological question, where the mean expression level can be obtained via previous studies or bulk RNA-Seq and is stable for cells within the same tissue (Supp. Fig. 1). We illustrate the proposed experimental design procedure by considering PBMCs with the corresponding marker genes (Fig. 1c). The goal is to ensure reliable estimation for all these genes that are above a certain expression level, say that of MS4A1. Hence, the expression level of the gene of interest, i.e. MS4A1, naturally defines the reliable detection limit *p\** at which we should guarantee an observation of one UMI per cell. Thus, given a budget B, choosing 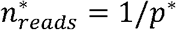 and 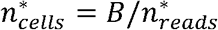 achieves the optimal trade-off for reliable detection at *p\**. In this example, MS4A1 will be sequenced ∼1 UMI per cell. Interestingly, this approach suggests a slightly deeper sequencing for current 10x datasets (Supp. Figs. 1-2). Moreover, although the exact optimal depth is task-dependent, our empirical evaluations have shown that the above recommendation is remarkably consistent across all quantities considered in this paper — typically lying in a narrow range between 0.2 and 1 (Fig. 1d, Supp. Fig. 4). Last but not the least, our trade-off analysis can also provide a post-hoc guidance for reliable estimation for existing datasets, namely for certain quantities, to determine which genes can be reliably estimated and which cannot, based on their mean expression level (Fig. 1e).

Another important result arising from our experimental design framework is the fundamental role of the estimator in the optimal trade-off. A very common — almost routine — practice in the literature is to use the so-called plug-in estimator, which, as a general recipe, blindly uses the read counts **Y**_1_, **Y**_2_,… as a proxy for the true gene expression levels **X**_1_, **X**_2_,… effectively estimating the corresponding distributional quantities by “plugging-in” the observed values. For example, the plug-in estimator naturally estimates the mean of the gene expression distribution *P*_**X**_ by that of *P*_**Y**_, the variance of *P*_**X**_ by that of *P*_**Y**_, etc. This approach, although very accurate for deeply sequenced datasets, becomes increasingly problematic in the limit of shallow sequencing; overdispersion and inflated dropout levels in lowly expressed genes, typically associated in the literature with scRNA-Seq, are some of the more pronounced consequences. For the sequencing budget allocation problem, we did not restrict our results to any particular estimator; our analysis suggested that the optimal trade-off cannot be achieved by the conventional plug-in approach but with another class of estimators developed via Empirical Bayes (EB, Supp. Note Table 1). EB estimators are inherently aware of the Poisson sampling noise introduced by sequencing, and therefore can adapt to varying sequencing depths. In Fig. 2 we provide a comprehensive evaluation of their performance in several key applications and show that they provide remarkably consistent estimates across varying sequencing depths and different datasets. Also, they are shown to be biologically meaningful (Fig. 2d, Supp. Fig 12). In contrast, the plug-in approach — being sensitive to the sequencing depth, significantly overestimates the variability in gene expression (cv) due to the inevitable zero-inflation occurring at shallow sequencing (Fig. 2a), and subsequently limits the performance of common downstream tasks such as PCA and gene network analysis (Methods, Fig. 2b-d).

## Methods

### Model

For a scRNA-Seq experiment, let *n*_*cells*_ be the number of cells and *n*_*reads*_ be the average UMIs per cell. The total number of UMIs *B* = *n*_*cells*_ × *n*_*reads*_ is used to denote the available budget for this experiment. Given a fixed budget, we are interested in the optimal allocation between *n*_*cells*_ and *n*_*reads*_ for estimating certain distributional quantities that are important to scRNA-Seq analysis.

We adopt an empirical Bayes model for the analysis (Supp. Note Sec. 1). Let *G* be the number of genes and for each cell *c* = 1, …, *n*_*cells*_, let **X**_*c*_=[*X*_*c*1_,…, *X*_*cG*_] ∈ *Δ*^*G*^ be the relative gene expression level, assumed to be drawn i.i.d. from some unknown cell distribution *P***_X_**. The gene expression level.**X**_***c***_is measured by the observed UMIs **Y**_*c*_∈ ℕ^*G*^ via sequencing, of which the stochastic process is modelled using Poisson noise. In addition, we assume a size factor *γ*_*c*_ for each cell that accounts for the variation in cell sizes. To summarize, the model can be written as

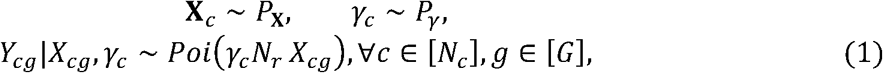

where *c* is the cell index and *g* is the gene index.

### Quantities to estimate

We consider five distributional quantities of P_**X**._ that are commonly-used in scRNA-Seq analysis, namely the moment, covariance matrix, inactive probability, pairwise inactive probability, and gene distribution (Supp. Note Sec. 2). The *k*-th gene moment is defined as *M*_*K,g*_ = 𝔼[*X*_*cg*_ ^*k*^] and the moments can be used to calculate quantities like mean expression level, coefficient of variation or Fano factor. The covariance matrix can be used for spectrum methods like PCA or spectral clustering. It also gives the Pearson correlation matrix, which can be used in gene network analysis. The inactive probability is defined as 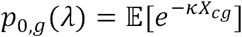 and is designed to quantify the proportion of cells where gene *g* is inactive. Since within a cell an inactive gene may still be measured at a low expression level, the exponential function is used to down-weight the small non-zero part of *P*_**X**_ The pairwise inactive probability, defined similarly as 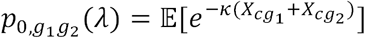, can further be used for gene co-expression analysis.

### Optimal budget allocation

We consider a single gene and derive the optimal budget allocation for estimating all the above quantities of its distribution 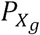 (Supp. Note Sec. 5). Since the mean relative expression level of a gene *P*_*g*_ is relatively stable within a specific tissue/sample and does not vary a lot across different experiments (Supp. Fig. 1), one can safely estimate that for an experiment with budget *B*, the total number of reads for gene S is around *p*_*g*_B. Then the trade-off with respect to gene *g* can be written as *p*_*g*_*B* = *n*_*reads,g*_ × *n*_*cells*_, where *n*_*reads,g*_ is the mean read counts per cell for gene *g*, satisfying the relation *n*_*reads,g*_ = *p*_*g*_*n*_*reads*_.

#### Theorem 1. Optimal budget allocation, informal

For estimating moments, covariance matrix, inactive probability, pairwise inactive probability and distribution, the optimal budget allocation is

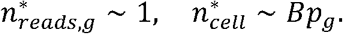

The optimality is in the sense of minimizing the worst-case error over a family of distributions 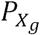 with mild assumptions and the optimal error rate is achieved by the EB estimators.

The expression 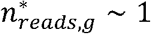 in Theorem 1 implies that the optimal sequencing depth is given by some constant independent of the sequencing budget. Therefore, for a scRNA-Seq experiment, we should aim at a certain sequencing depth; when the budget increases, we should keep the same depth and allocate the additional budget towards collecting more cells. In other words, after having achieved a certain sequencing depth, deeper sequencing does not help as much as having more cells. We also note that the actual value of this optimal sequencing depth may be different for estimating different quantities, which is further investigated in the following section. In addition, Theorem 1 suggests that EB estimators should be used for optimal estimation, whose effectiveness is demonstrated in Fig. 2.

### Experimental design

The exact values of the optimal sequencing depth 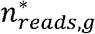 for estimating different quantities are investigated both theoretically and via simulations. First, the closed-form expressions of the optimal depth 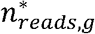are derived for estimating the mean, the second moment, and the gamma parameters, which depend on the distribution 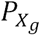 but are nonetheless around 1 for typical cases (Supp. Note Lemma 2). Second, estimation errors under different budget splits are simulated by subsampling from an existing dataset with deeply sequenced genes and many cells (top 72 genes of brain_1.3m, Fig. 1d), where the subsample procedure is provable to closely resemble the reality (Supp. Note. 6.4). Third, a more controlled simulation that assumes the Poisson model is conducted to provide a more comprehensive evaluation (Supp. Fig. 2). Both simulations exhibit similar qualitative behaviours and imply that the optimal sequencing depth 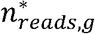 for estimating different quantities are between 0.2 and 2. Therefore, we reach the conclusion that the optimal budget allocation for a single gene is to have ∼1 read per cell.

When there are many genes of interest, the gene among them with the smallest relative mean expression level becomes the bottleneck since it has the least number of reads (Fig. 1c, top). We call its relative mean expression level *p\** the reliable detection limit, below which the estimation performance cannot be guaranteed. The optimal sequencing depth for the entire experiment 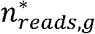 is chosen so that the gene at the reliable detection limit has 1 read per cell, which minimizes the worst-case error for all genes of interest. Compared to this optimal allocation, deeper sequencing (green) gives a homogeneous error across genes but at a much higher level, while a shallower sequencing (blue) gives a small error for a few highly expressed genes but its performance quickly deteriorates (Fig. 1c, bottom).

The recommended budget allocation in general suggests a slightly deeper sequencing depth as compared to existing datasets, e.g. 7k UMIs per cell for the pbmc_4k dataset considering MS4A1 and 14k UMIs per cell for the brain_9k dataset considering S100a10 (Fig. 1b, Supp. Fig. 2). Such a depth is feasible for the current 10x Genomics technology, which is estimated to be able to sequence 10-45k UMIs per cell where the actual values depend on different tissues (Supp. Note Sec. 3.1). In addition, under such sequencing depth all analyses are valid since the Poisson model is still a good approximation of the sequencing process. Regarding the rare genes, since the UMI efficiency for the 10x technology is estimated to be 10-15%, in order to achieve one read per cell, the gene needs to have at least 1/0.15∼7 transcripts in the cell. The gene CD4 (Supp. Fig. 2b) seems to be below this limit. For such genes, the recommendation should be sequencing until saturation.

### Designing EB estimators

The empirical Bayes (EB) estimators refer to the estimators that are aware of the noise model (which is Poisson here) and correct for the noise introduced by it. As an illustrating example, consider a simplified model:

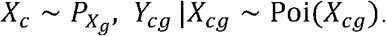

The plug-in estimator estimates the gene variance by the sample variance of UMIs, i.e.

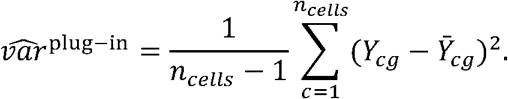

The problem with this is that the estimated value is usually overly variable due to the presence of the Poisson noise. Indeed, a simple calculation shows that

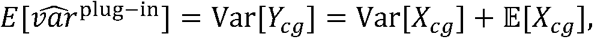

where the second term 𝔼[*X*_*cg*_] corresponds to the technical variation introduced by the Poisson noise. Then conceptually we can write:

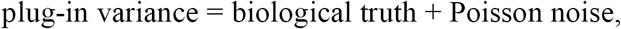

from which we can see that the plug-in estimate is overly inflated by the Poisson noise. In this case, this bias can be easily corrected by simply subtracting the mean; the corresponding EB variance estimator can be written as

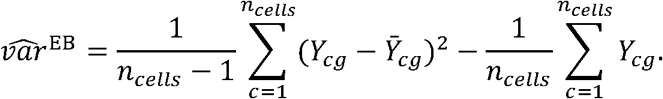

For an extensive comparison of plug-in and EB estimators in various cases, see Supp. Note, Table 1 and Sec. 4.

### Empirical evaluation of the trade-off

We conducted two sets of simulations to evaluate the estimation error under different budget splits, which differ in how the data are generated. The first simulation (Fig. 1d) subsamples from a high-budget dataset consisting of the top 72 genes from the brain_1.3m dataset, where each gene has at least 10 reads per cell and there are 1.3 million cells. This simulation better resembles the reality as the subsample procedure does not assume the Poisson model (Supp. Note 6.4). However, since we do not know the true gene distribution, the plug-in estimates of the high-budget dataset that we subsample from are used as proxies against which we evaluate the estimation error. The second simulation generates the data according to model (1), where the true gene distribution *P*_**X**_ is obtained by using the empirical distribution of the first 100 highly-expressed genes in the pbmc_4k dataset (Supp. Fig. 4). This setting better validates the theory since it assumes the same model. Moreover, the estimation error is exact since the ground truth is available. Both simulations include many genes to address the heterogeneity of the gene distribution and the genes considered here, being top genes in the dataset, have similar mean expression levels so that the mean reads over all genes can well represent the mean reads for each gene. Both simulations exhibit similar qualitative behaviours that the optimal depth (mean reads per cell per gene) is a constant that does not depend on the budget, implying the theory matches the reality.

### Post-hoc guidance for reliable estimation

The feasible region (top) and the post-hoc table (bottom) are obtained via simulation, where we fix the number of cells (1k, 5k, 10k, 30k, 70k) and investigate how the error decreases as a function of the depth (Supp. Fig. 5-6). The data are generated according to model (1) similar to the second trade-off simulation, where the empirical distributions of the marker genes in pbmc_4k and brain_9k are used as the true gene distribution respectively to account for heterogeneity in different tissues. The true gene distribution is normalized so that each gene has the same mean expression level. As a result, the mean reads over all genes is exactly the mean reads for each gene, providing a single-gene level error characterization. The post-hoc table is obtained by finding the smallest depth such that the relative error is smaller than 0.1(−2 in the log10 scale for the relative squared error and -1 for other errors, Supp. Sec. 6.3). The results for both simulations are very similar. Hence only the table for pbmc_4k is included.

### Comparing the performance of plug-in and EB estimators

Fig. 2a demonstrates that the EB estimator is adaptive to different sequencing depth while the plug-in estimator is not. The top panel shows the estimated cv using plug-in and EB under different sequencing depths, where we can see a clear inflation for plug-in. The full data is from pbmc_4k and the subsample rate ranges from 0.2 to 1 (full data). The experiment is repeated 5 times and the 3-std confidence interval is provided. The results for other genes as well as for estimating the inactive probability can be found in Supp. Fig. 7. The middle panel compares the estimated cv from two datasets of the same tissue. Genes with at least 0.1 reads per cell are considered since we believe genes below this level cannot be reliably estimated. The EB estimator may produce an invalid result when the plug-in variances are smaller than the plug-in means, where the Poisson model will break down. Such cases are not common and are excluded while counting the number of genes above/below the red line. Hence the total number genes for the two panels may slightly differ. More results are in Supp. Fig. 8-9. The bottom panel shows that EB can recover the gene distribution from a shallow sequencing data. The shallow data is generated by subsampling from the full data and is 5times shallower. To evaluate the error, the recovered distribution is rescaled to have the same mean as the empirical distribution from the full data. More results are in Supp. Fig. 10.

Fig. 2b investigates a common task that the features (genes) are selected based on cv and PCA is then performed on selected features. The data is from pbmc_4k and is clipped at the 99th quantile to prevent outliers. Such procedure also appears in previous works on PCA [11]. The top 500 genes with the highest cv are selected and the PCA scores are plotted for the 2nd and 3rd PC direction. The first direction is skipped because it usually corresponds to the variation in cell sizes. The results on the full data and a subsampled data (3 times shallower) are compared, where EB is more consistent than plug-in.

Fig. 3c considers recovering gene functional group using Pearson correlation. We use the pbmc_4k dataset here since the biological structure of the PBMCs is well-understood. The major cell populations identified in this dataset are T-cells (IL7R, CD3D/E/, LCK), NK-cells (NKG7, PRF1, KLRD1, GZMA, HOPX, CST7), B-cells (CD79A, BANK1, IGHD, LINC00926, MS4A1), myeloid-derived cells (S100A8/9, MNDA, FGL2, CLEC7A, IFI30) and megakaryocytes/platelets (PF4, PPBP). The heatmap of the EB estimated Pearson correlation of those genes are visualized in Fig. 3c top, which shows that EB can well capture the gene functional groups. A subsample experiment is then conducted to investigate how well the estimators can recover the modules from the shallow sequencing data. The data are subsampled from the full data with the rate 100% (full), 25%, 10% and 5%. EB can recover the module at a much shallower depth as compared to plug-in.

### Gene network analysis of the 10x Genomics’ pbmc_4k dataset

The gene network (Fig. 2d top) is constructed based on the EB estimated Pearson correlation using the pbmc_4k dataset. The genes are filtered based on whether the EB estimated variance is larger than 0.1, which leaves 791 genes. A correlation larger than 0.8 is considered as a gene-gene edge. We found that varying the threshold from 0.4 to 0.95 does not significantly affect the result. The gene modules are identified based on knowledge of marker genes and gene pathways as well as previous studies on PBMCs (Supp. Note 6.5). We also note that the existence of megakaryocytes/platelets may be due to the imperfection of PBMC isolation and since many genes are expressed in multiple cell populations (e.g. CD74, CD27), the resulting annotation only gives a rough picture of the underlying gene functional groups.

Next, we consider some important genes and plot their correlations with all other genes (Fig. 2d middle, Supp. Fig. 11). As a general phenomenon, the EB estimated values are more spread out and exhibit different modes corresponding to genes that interact differently with the gene of interest. The plug-in estimated values are nonetheless much closer to zero even for the genes that are known to be well-correlated.

Finally, we consider the gene pairs where the estimated values for EB and plug-in differ significantly (>0.7). Out of 1054 such pairs, 91 are also annotated based on STRING [20], yielding a p-value of 4.2e-11 while testing against the null that these pairs are selected at random (Supp. Note 6.5). We plot the histograms of several such pairs and show that all of them have clear biological interpretations (Fig. 2d bottom, Supp. Fig. 12). LY86 (also known as MD1) is a secreted protein that has been shown to play an important role in T-cell activation while CD3E is expressed within T-cells (Supp. Note 6.5). These two genes are not co-expressed and hence are negatively correlated. POMP encodes a chaperone for proteasome assembly while PSMA7 is one of the 17 essential subunits for the complete assembly of the 20S proteasome complex. Hence, the two genes work together for proteasome assembly and should be positively correlated. In spite of the strong biological evidence, the plug-in estimator produces very small values due to the presence of sequencing noise (See also Supp. Fig. 12).

## Supplementary Information

supplementary_figures.pdf contains additional figures compensating the experiments in the main paper. supplementary_note.pdf contains the theoretical analysis as well as more technical details.

## Acknowledgments

This research was in part motivated by discussions on the experimental design question in the Human Cell Atlas First Annual Jamboree meeting. We would like to thank Lior Pachter for his valuable input and constructive suggestions throughout the course of this study; Jase Gehring, Wenying Pan, and Taibo Li for their helpful feedback. Thanks also to Patrick Marks for very useful feedback on an earlier version of the paper. MZ is partially supported by Stanford Graduate Fellowship.

## Author Contributions

All authors contributed extensively to the work presented in this paper.

## Competing Interests Statement

The authors declare that they have no competing financial interests.

## Correspondence

Requests for materials should be addressed to DT (e-mail: dntse*@*stanford.edu).

## Code Availability

We developed the python package *sceb* (single-cell empirical Bayes) for the EB estimators used in this paper (available on PyPI). The code to reproduce all experiments and generate the figures presented in this paper can be found at https://github.com/martinjzhang/single_cell_eb

## Data Availability

The datasets that we use are were generated by 10x Genomics’ v2 chemistry [17]. pbmc_4k, pbmc_8k contain peripheral blood mononuclear cells (PBMCs) from a healthy donor (the same donor). brain_1k, brain_2k, brain_9k, brain_1.3m contain cells from a combined cortex, hippocampus and sub ventricular zone of an E18 mouse. The pair 293T_1k, 3T3_1k contain 1:1 mixture of fresh frozen human (HEK293T) and mouse (NIH3T3) cells. So are the pairs 293T_6k, 3T3_6k and 293T_12k, 3T3_12k. The links of the datasets:

pbmc_4k:https://support.10xgenomics.com/single-cell-gene-expression/datasets/2.1.0/pbmc4k

pbmc_8k:https://support.10xgenomics.com/single-cell-gene-expression/datasets/2.1.0/pbmc8k

brain_1k:https://support.10xgenomics.com/single-cell-gene-expression/datasets/2.1.0/neurons_900

brain_2k:https://support.10xgenomics.com/single-cell-gene-expression/datasets/2.1.0/neurons_2000

brain_9k:https://support.10xgenomics.com/single-cell-gene-expression/datasets/2.1.0/neuron_9k

brain_1.3m:https://support.10xgenomics.com/single-cell-gene-expression/datasets/1.3.0/1M_neurons

293T_1k,3T3_1k:https://support.10xgenomics.com/single-cell-gene-expression/datasets/2.1.0/hgmm_1k

93T_6k,3T3_6k:https://support.10xgenomics.com/single-cell-gene-expression/datasets/2.1.0/hgmm_6k

293T_12k,3T3_12k: https://support.10xgenomics.com/single-cell-gene-expression/datasets/2.1.0/hgmm_12k

